## Supplements for "Executive Resources Shape the Impact of Language Predictability Across the Adult Lifespan"

* Shared first authorship

★ Shared last authorship

### Supplementary Methods

#### Demographics

We recruited participants both using the participant database of the Max-Planck-Institute in Leipzig and the online participant recruitment platform *Prolific*. Due to the predominantly younger demographic on *Prolific*, the online samples comprised mainly younger and middle-aged participants, whereas the lab sample spanned a broader age range (see Fig. S1).

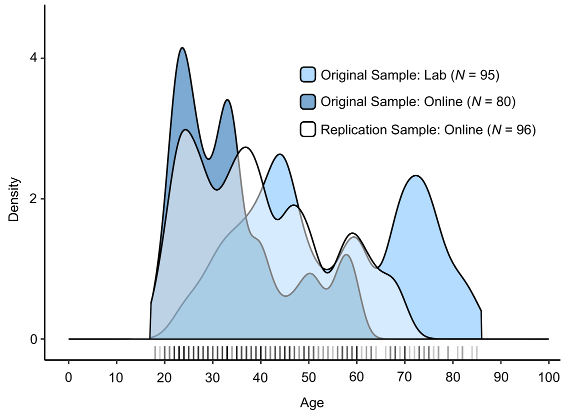


**Figure S1. Comparison of age distribution between samples.**

### Study Design

#### *Text stimuli*

During the experiment, participants were asked to read short newspaper articles on emotionally neutral topics such as literature, history, geography and biology. All texts were edited to be easy to understand without excessive simplification. By doing so, we aimed to keep the cognitive load in the reading task as low as possible, while still maintaining a balance between text clarity and the avoidance of consistently high word predictability.

All texts had a Wiener Sachtextformel ($WSTF_4$; [118]) score below or equal to 10, which corresponds to a reading level suitable for students below 10th grade (mean $WSTF_4$ score: 7.9 ± 0.623). As the $WSTF_4$ only measures the syntactic complexity of a text, participants rated the subjective text difficulty as well as their subjective interest for two of the texts presented to them. All texts used in this study yielded a mean difficulty rating of 21.983 ± 2.799 (on a scale from 0 to 100, with 100 being "extremely difficult") and a mean interest rating of 70.684 ± 5.657 (on a scale from 0 to 100, with 100 being "extremely interesting"), confirming that the texts used in this study were both easy to understand and interesting to read.

#### *Structure of the Experiment*

At the outset of the experiment, participants were first presented with a training block of the Reading Only condition to familiarise themselves with the task. This was followed by the first main block of the Reading Only task (300 trials). After this, in a short training block (20 trials, repeating it was optional) followed by a longer main block (60 trials in the online experiment, 90 trials in the lab experiment), the participant was introduced to either the 1-back or the 2-back task as a non-linguistic single task comprising coloured rectangles as stimuli. Which n-back task was introduced first was randomised. This was then followed by the first dual-task block (300 trials) where the previously practised n-back task was performed together with the reading task. After having completed the first dual-task block in one of the two n-back conditions, the participant was then introduced to the other n-back task in the same fashion as before. Having performed each of the conditions once, the participant was subsequently presented with three main blocks of each of the three conditions in random order (300 trials each). After each block comprising a reading task, participants were asked to answer three multiple-choice comprehension questions on the content of the text.

### Supplementary Results

#### Table S1. Results from models for task performance measures (N = 175).

| | LMM for d-primes | | | | | GLMM for comprehension question accuracy | | | |
|---|---|---|---|---|---|---|---|---|---|
| | *Estimate* | *Std. Error* | *t* | *df* | *p* | *OR* | *Std. Error* | *z* | *p* |
| mean d-prime single tasks | 0.469 | 0.0549 | 8.550 | 166.51 | $2.294 \times 10^{-14}$ * | | | | |
| mean comprehension question performance | 0.011 | 0.0036 | 3.053 | 171.653 | $3.943 \times 10^{-3}$ * | | | | |
| de-meaned comprehension question performance | -0.001 | 0.0012 | -0.475 | 372.688 | $6.350 \times 10^{-1}$ | | | | |
| block number | -0.006 | 0.0085 | -0.676 | 349.333 | $5.618 \times 10^{-1}$ | | | | |
| recording location [online] | -0.506 | 0.0902 | -5.609 | 163.182 | $1.920 \times 10^{-7}$ * | 0.980 | 0.1876 | -0.106 | $9.155 \times 10^{-1}$ |
| age | -0.005 | 0.0027 | -2.057 | 164.038 | $5.003 \times 10^{-2}$ | 0.986 | 0.0053 | -2.676 | $1.304 \times 10^{-2}$ * |
| cognitive load [1-back vs. Reading Only] | | | | | | 0.253 | 0.0390 | -8.928 | $1.011 \times 10^{-18}$ * |
| cognitive load [2-back vs. Reading Only] | | | | | | 0.156 | 0.0234 | -12.403 | $1.753 \times 10^{-34}$ * |
| cognitive load. [2-back vs. 1-back] | -1.636 | 0.0626 | -26.120 | 173.125 | $2.672 \times 10^{-61}$ * | | | | |
| age * cognitive load [1-back vs. Reading Only] | | | | | | 0.990 | 0.0081 | -1.183 | $3.313 \times 10^{-1}$ |
| age * cognitive load [2-back vs. Reading Only] | | | | | | 1.003 | 0.0080 | 0.395 | $8.081 \times 10^{-1}$ |
| age * cognitive load [2-back vs. 1-back] | -0.014 | 0.0035 | -3.931 | 169.766 | $2.210 \times 10^{-4}$ * | | | | |
| model fit | *Conditional / Marginal $R^2$* | | | *ICC* | | *Conditional / Marginal $R^2$* | | *ICC* | |
| | 0.822 / 0.634 | | | 0.512 | | 0.304 / 0.146 | | 0.185 | |

*Note*. All continuous predictors were centred. Degrees of freedom for *p*-values, standard errors and confidence intervals (CI) were computed using Satterthwaite's approximation (LMM for d-primes) and Wald's approximation (GLMM for comprehension question accuracy). All *p*-values reported here are FDR-corrected and were computed using ANOVAs with type III sums of squares. Results that are significant on an alpha-level of 0.05 are marked with a star. OR = Odds Ratio.

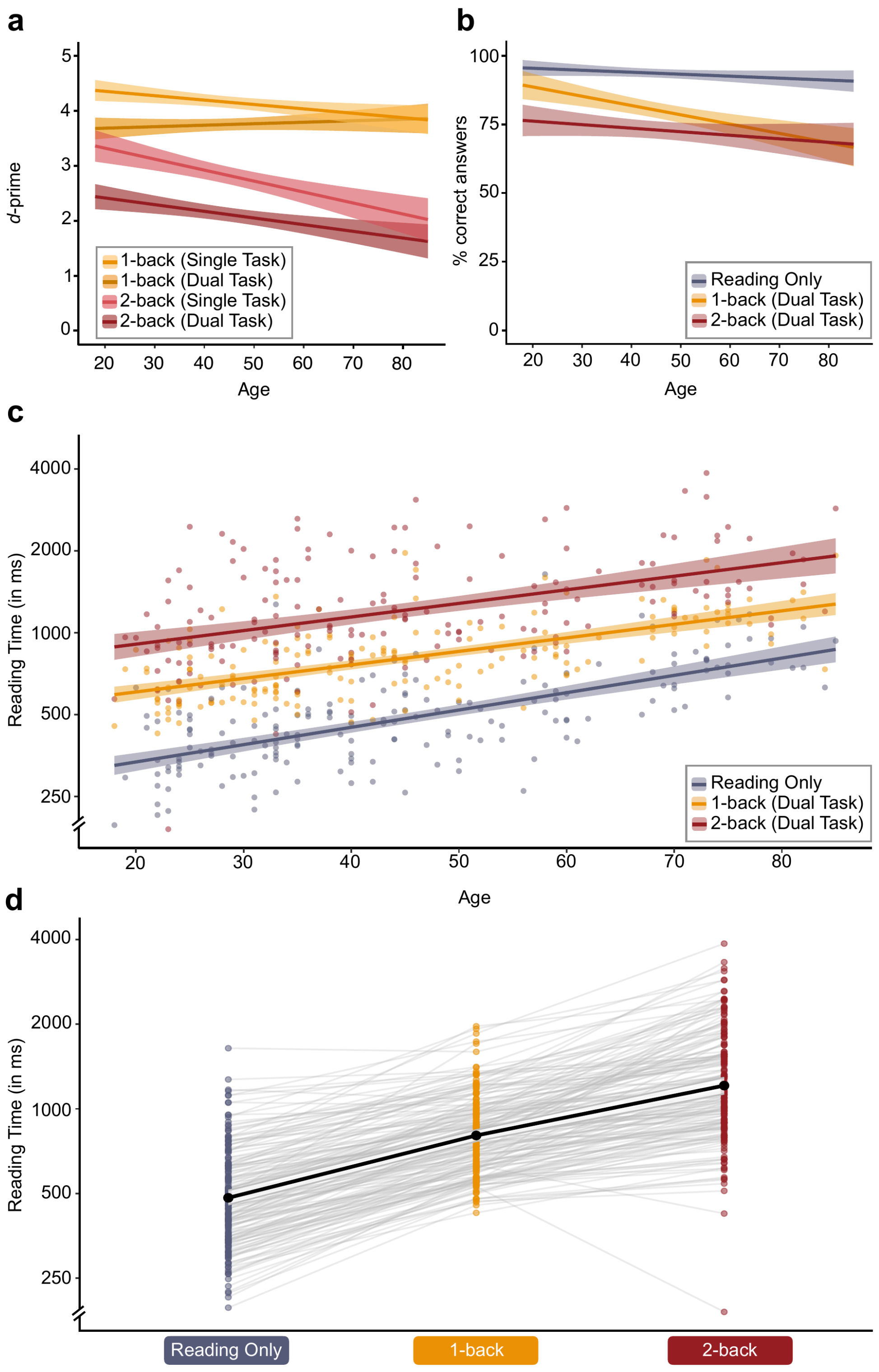


**Figure S2. Task performance and reading times by age and cognitive load condition.** Task performance (d-primes) in conditions with an n-back task (**a**). Accuracy in the comprehension question performance task (**b**). Reading Times by age and condition (**c**). Reading times by condition (**d**). Solid line: M, shaded area: 95% CI, point: mean reading time for one participant in the respective condition.

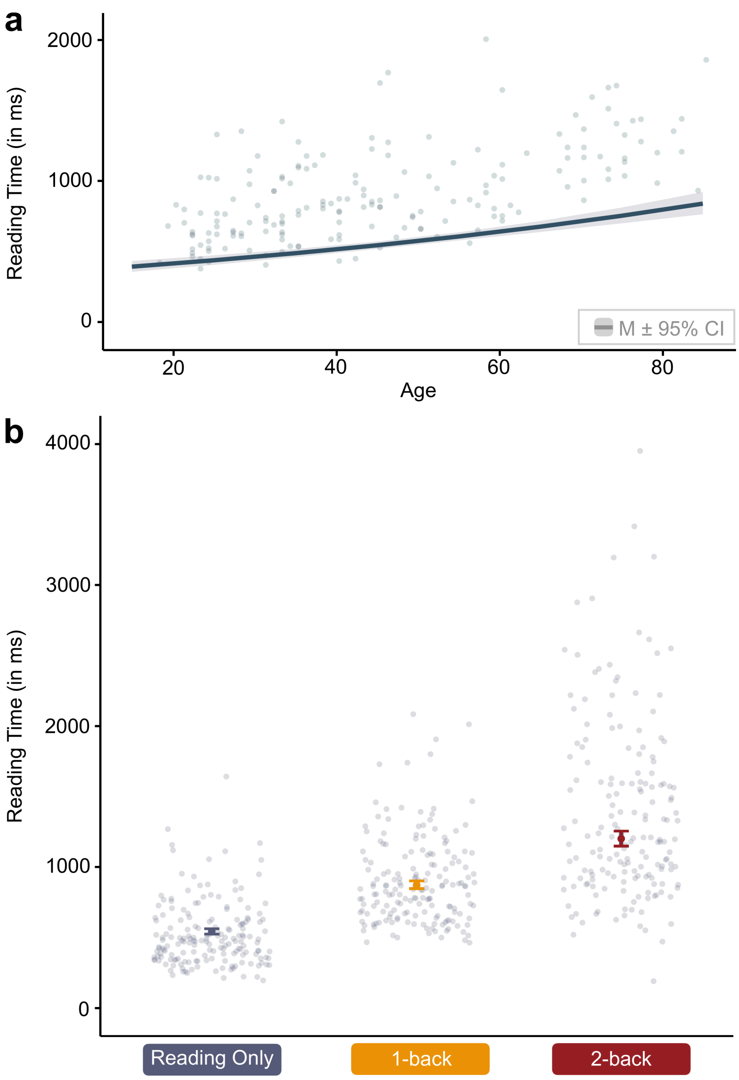


**Figure S3. Individual predicted reading times.** Reading times by age (**a**) and reading times by condition (**b**).

**Figure S4. Comparison of factor smooths for different levels of cognitive load from the three-way interaction of age, surprisal, and cognitive load.** The difference smooths show slightly stronger

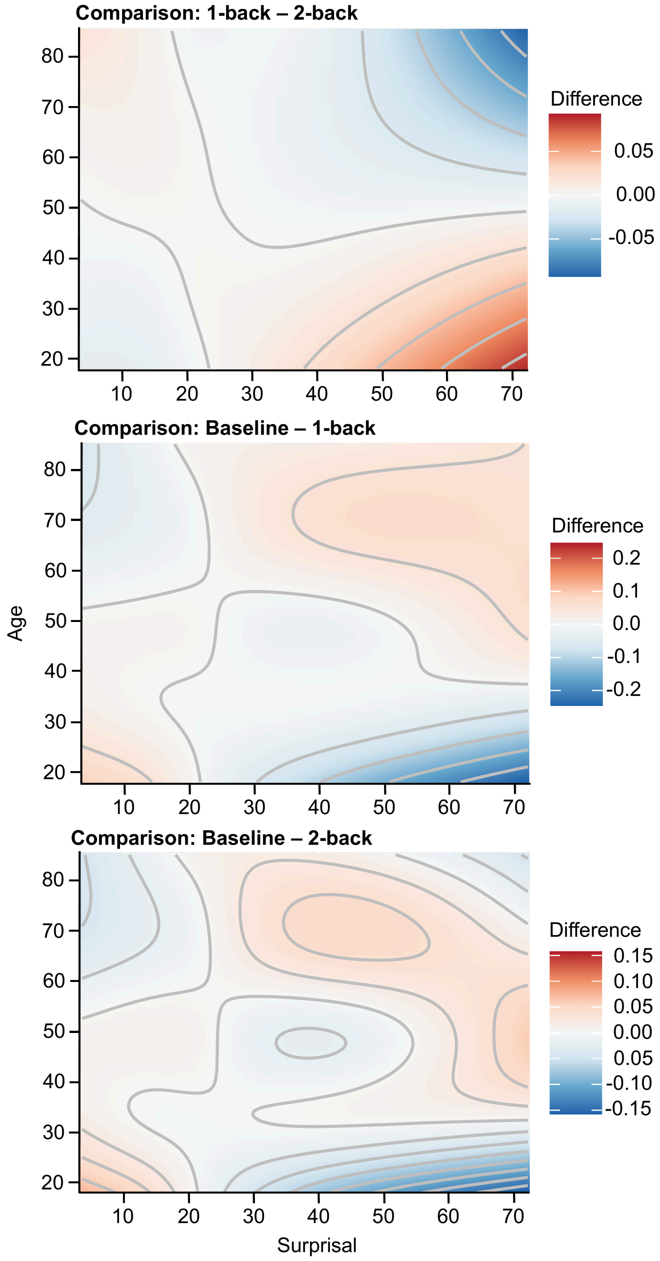


effects of high surprisal in young than older adults for the 1-back relative to the 2-back condition, and stronger effects of high surprisal in older adults for the Reading only relative to the n-back condition.

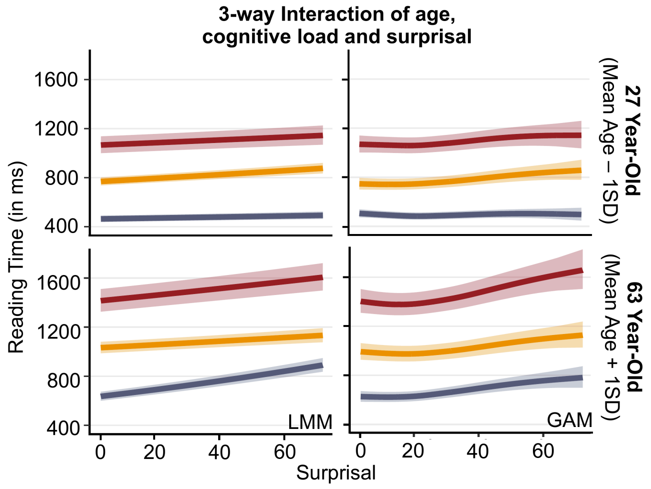


**Figure S5. Comparison of the results of the LMM and GAM control analyses.** Panel **c** illustrates the interaction between cognitive load and surprisal for a representative younger and older participant, estimated using the LMM (left) and the GAM (right). For a complementary visualisation of the three-way interaction between age, cognitive load, and surprisal, see Fig. S4. For a visualisation of the main effects of age, surprisal and cognitive load on reading time, please see Fig. 3c.

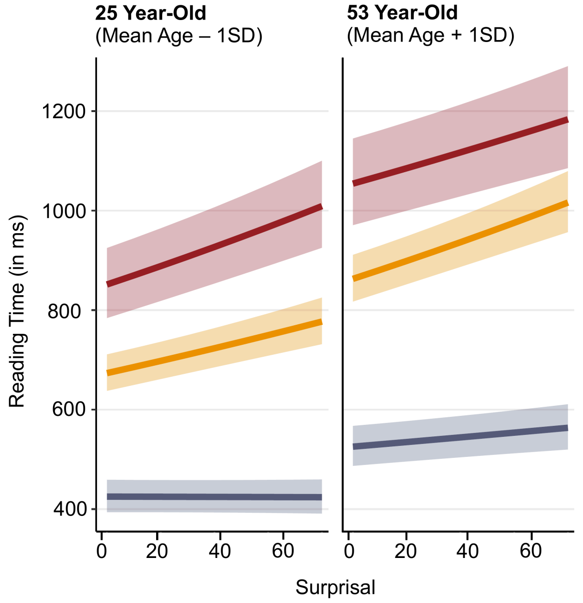


**Figure S6. Three-way interaction of age, surprisal, and cognitive load in the replication sample.**

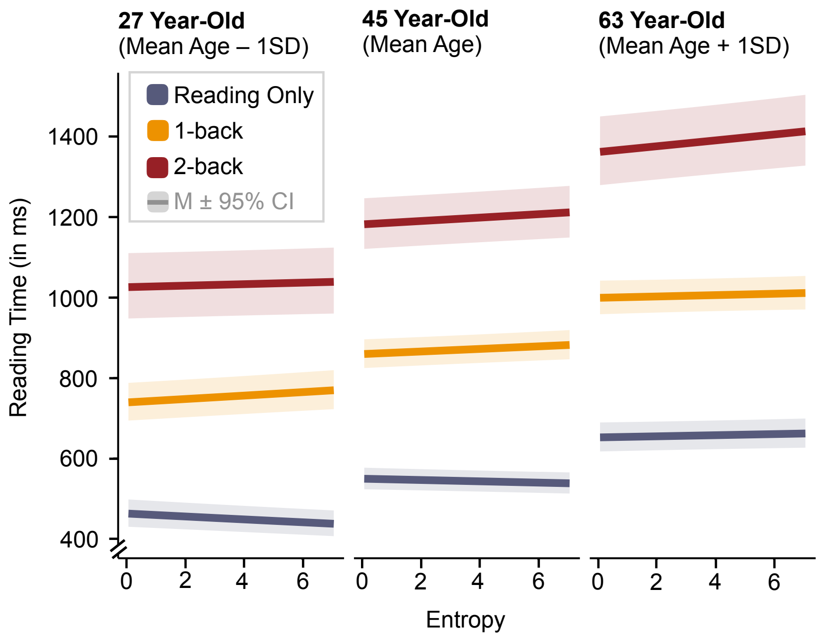


**Figure S7. Three-way interaction of age, entropy, and cognitive load in the full sample (N = 175).**

**Table S2. Results from the model for reading times for full original sample (N = 175).**

| | | LMM for Full Original Sample (N = 175) | | | | | |
|---|---|---|---|---|---|---|---|
| | ***Predictors*** | ***Estimate*** | ***Std. Error*** | ***CI*** | ***t*** | ***df*** | ***p*** |
| **Main Effects** | reading time of previous trial (log-transformed) | 0.110235 | 0.001409 | 0.107472 – 0.112997 | 78.210829 | 287711.33 | $< 1.33 \times 10^{-322}$ * |
| | d-prime | -0.006293 | 0.001767 | -0.009755 – -0.002831 | -3.562277 | 224716.70 | $4.903 \times 10^{-4}$ * |
| | mean d-prime singletasks | 0.084636 | 0.019585 | 0.045974 – 0.123298 | 4.321418 | 169.86 | $3.717 \times 10^{-5}$ * |
| | mean comprehension question performance | 0.002841 | 0.001276 | 0.000321 – 0.005360 | 2.225728 | 169.26 | $3.282 \times 10^{-2}$ * |
| | de-meaned comprehension question performance | 0.000053 | 0.000038 | -0.000023 – 0.000128 | 1.374475 | 257871.34 | $1.693 \times 10^{-1}$ |
| | word frequency | 0.643570 | 0.362057 | -0.067231 – 1.354371 | 1.777538 | 727.78 | $8.280 \times 10^{-2}$ |
| | word length | 0.007839 | 0.000407 | 0.007039 – 0.008638 | 19.240951 | 1400.80 | $7.407 \times 10^{-73}$ * |
| | word entropy | 0.001410 | 0.000785 | -0.000129 – 0.002949 | 1.796533 | 7594.27 | $8.280 \times 10^{-2}$ |
| | n-back reaction [reaction vs. no reaction] | 0.317464 | 0.001799 | 0.313937 – 0.320990 | 176.444743 | 287862.05 | $< 1.33 \times 10^{-322}$ * |
| | block number | -0.006495 | 0.000164 | -0.006816 – -0.006173 | -39.637620 | 287253.02 | $< 1.33 \times 10^{-322}$ * |
| | trial number | -0.000456 | 0.000007 | -0.000470 – -0.000442 | -64.085235 | 18673.68 | $< 1.33 \times 10^{-322}$ * |
| | recording location [online vs. lab] | -0.219339 | 0.032284 | -0.283072 – -0.155606 | -6.793975 | 168.69 | $2.686 \times 10^{-10}$ * |
| | surprisal | 0.001707 | 0.000151 | 0.001411 – 0.002002 | 11.320677 | 2361.37 | $1.368 \times 10^{-28}$ * |
| | Age | 0.009113 | 0.000991 | 0.007158 – 0.011068 | 9.199100 | 178.46 | $1.751 \times 10^{-16}$ * |
| | cognitive load [1-back vs. Reading Only] | 0.473800 | 0.013916 | 0.446336 – 0.501264 | 34.046321 | 176.18 | $8.399 \times 10^{-79}$ * |
| | cognitive load [2-back vs. Reading Only] | 0.791540 | 0.026090 | 0.740046 – 0.843034 | 30.338989 | 173.76 | $7.320 \times 10^{-71}$ * |
| **2-way Interactions** | surprisal x age | 0.000035 | 0.000004 | 0.000027 – 0.000042 | 9.287151 | 287771.27 | $3.481 \times 10^{-20}$ * |
| | surprisal x cognitive load [1-back vs. Reading Only] | -0.001093 | 0.000161 | -0.001409 – -0.000776 | -6.771521 | 287959.11 | $2.043 \times 10^{-11}$ * |
| | surprisal x cognitive load [2-back vs. Reading Only] | -0.001255 | 0.000163 | -0.001575 – -0.000935 | -7.681261 | 288294.96 | $2.709 \times 10^{-14}$ * |
| | age x cognitive load [1-back vs. Reading Only] | -0.002798 | 0.000776 | -0.004330 – -0.001267 | -3.606479 | 171.99 | $5.135 \times 10^{-4}$ * |
| | age x cognitive load [2-back vs. Reading Only] | -0.002458 | 0.001454 | -0.005329 – 0.000412 | -1.690400 | 170.79 | $9.681 \times 10^{-2}$ |
| **3-way Interactions** | surprisal x age x cognitive load [1-back vs. Reading Only] | -0.000111 | 0.000009 | -0.000129 – -0.000094 | -12.266076 | 287807.34 | $3.748 \times 10^{-34}$ * |
| | surprisal x age x cognitive load [2-back vs. Reading Only] | -0.000078 | 0.000009 | -0.000096 – -0.000060 | -8.483676 | 287771.65 | $4.384 \times 10^{-17}$ * |
| **Model Fit** | Intra-class Correlation (ICC) | 0.46 | | | | | |
| | Marginal $R^2$ / Conditional $R^2$ | 0.643 / 0.807 | | | | | |

*Note.* All continuous predictors were centred. Degrees of freedom for *p*-values, standard errors and confidence intervals (CI) were computed using Satterthwaite's approximation. All *p*-values reported here are FDR-corrected and were computed using ANOVAs with type III sum of squares. Results that are significant on an alpha-level of 0.05 are marked with a star.

**Table S3. Results from models for reading times for original online sample and online replication sample (N = 80 and N = 96, respectively).**

| | | LMM for Online Original Sample (N = 80) | | | | | | LMM for Online Replication Sample (N = 96) | | | | | |
|---|---|---|---|---|---|---|---|---|---|---|---|---|---|
| | ***Predictors*** | ***Estimate*** | ***Std. Error*** | ***CI*** | ***t*** | ***df*** | ***p*** | ***Estimate*** | ***Std. Error*** | ***CI*** | ***t*** | ***df*** | ***p*** |
| **Main Effects** | reading time of previous trial (log-transformed) | 0.076157 | 0.002060 | 0.072120 – 0.080193 | 36.978112 | 133340.940 | **$5.049 \times 10^{-297}$** * | 0.149035 | 0.001818 | 0.145472 – 0.152597 | 81.985 | 161495.845 | **< $3.442 \times 10^{-281}$** * |
| | d-prime | 0.058086 | 0.003309 | 0.051601 – 0.064571 | 17.555499 | 58135.052 | **$2.440 \times 10^{-68}$** * | -0.010767 | 0.002175 | -0.015029 – -0.006504 | -4.950607 | 139351.725 | **$8.888 \times 10^{-7}$** * |
| | mean d-prime singletasks | 0.093533 | 0.027543 | 0.038674 – 0.148391 | 3.395825 | 75.930447 | **$1.403 \times 10^{-3}$** * | 0.111742 | 0.020097 | 0.071832 – 0.151652 | 5.560231 | 92.614 | **$3.324 \times 10^{-7}$** * |
| | mean comprehension question performance | 0.003366 | 0.001733 | -0.000085 – 0.006816 | 1.942677 | 76.169 | $6.690 \times 10^{-2}$ | 0.003459 | 0.001659 | 0.000163 – 0.006754 | 2.084567 | 91.938 | **$4.487 \times 10^{-2}$** * |
| | de-meaned comprehension question performance | -0.000481 | 0.000059 | -0.000597 – -0.000365 | -8.123202 | 118882.498 | **$8.250 \times 10^{-16}$** * | -0.000715 | 0.000047 | -0.000807 – -0.000624 | -15.286171 | 152148.746 | **$2.661 \times 10^{-52}$** * |
| | word frequency | 0.255248 | 0.300091 | -0.335305 – 0.845800 | 0.850567 | 299.741 | $4.190 \times 10^{-1}$ | 0.277510 | 0.253935 | -0.222530 – 0.777550 | 1.092838 | 258.972 | $2.917 \times 10^{-1}$ |
| | word length | 0.006329 | 0.000414 | 0.005517 – 0.007141 | 15.297105 | 1292.834 | **$2.874 \times 10^{-48}$** * | 0.006341 | 0.000362 | 0.005631 – 0.007052 | 17.512071 | 1287.953 | **$2.794 \times 10^{-61}$** * |
| | word entropy | -0.000656 | 0.000933 | -0.002485 – 0.001173 | -0.702998 | 3838.651 | $4.821 \times 10^{-1}$ | 0.000337 | 0.000826 | -0.001283 – 0.001957 | 0.407433 | 3601.355 | $6.837 \times 10^{-1}$ |
| | n-back reaction [reaction vs. no reaction] | 0.366351 | 0.002603 | 0.361250 – 0.371452 | 140.757916 | 133345.032 | **< $5.049 \times 10^{-297}$** * | 0.335449 | 0.002293 | 0.330955 – 0.339943 | 146.303524 | 161785.798 | **< $3.442 \times 10^{-281}$** * |
| | block number | -0.008486 | 0.000247 | -0.008970 – -0.008002 | -34.367684 | 131940.893 | **$4.797 \times 10^{-257}$** * | -0.008042 | 0.000224 | -0.008481 – -0.007604 | -35.946825 | 159855.434 | **$3.442 \times 10^{-281}$** * |
| | trial number | -0.000407 | 0.000009 | -0.000425 – -0.000390 | -45.720722 | 8566.066 | **< $5.049 \times 10^{-297}$** * | -0.000386 | 0.000008 | -0.000402 – -0.000371 | -48.601047 | 8175.168 | **< $3.442 \times 10^{-281}$** * |
| | surprisal | 0.001145 | 0.000162 | 0.000826 – 0.001463 | 7.046510 | 1889.625 | **$4.190 \times 10^{-12}$** * | 0.001375 | 0.000144 | 0.001093 – 0.001656 | 9.578258 | 1886.753 | **$5.358 \times 10^{-21}$** * |
| | age | 0.005382 | 0.001535 | 0.002326 – 0.008438 | 3.507190 | 76.056 | **$1.057 \times 10^{-3}$** * | 0.008953 | 0.001318 | 0.006336 – 0.011570 | 6.795181 | 91.979 | **$1.460 \times 10^{-9}$** * |
| | cognitive load [1-back vs. Reading Only] | 0.468989 | 0.018722 | 0.431749 – 0.506229 | 25.050620 | 82.526 | **$5.650 \times 10^{-40}$** * | 0.507557 | 0.022113 | 0.463669 – 0.551446 | 22.953170 | 96.841 | **$1.343 \times 10^{-40}$** * |
| | cognitive load [2-back vs. Reading Only] | 0.824086 | 0.034653 | 0.755102 – 0.893071 | 23.781266 | 78.282 | **$2.909 \times 10^{-37}$** * | 0.722423 | 0.031793 | 0.659316 – 0.785531 | 22.722673 | 96.183 | **$3.806 \times 10^{-40}$** * |
| **2-way Interactions** | surprisal x cognitive load [1-back vs. Reading Only] | 0.000786 | 0.000223 | 0.000349 – 0.001223 | 3.522647 | 133382.112 | **$6.411 \times 10^{-4}$** * | 0.001499 | 0.000203 | 0.001101 – 0.001897 | 7.376731 | 161262.305 | **$2.667 \times 10^{-13}$** * |
| | surprisal x cognitive load [2-back vs. Reading Only] | 0.000375 | 0.000225 | -0.000067 – 0.000816 | 1.661967 | 133507.349 | $1.086 \times 10^{-1}$ | 0.001365 | 0.000203 | 0.000967 – 0.001763 | 6.721136 | 161923.055 | **$2.714 \times 10^{-11}$** * |
| **Model Fit** | Intra-class Correlation (ICC) | | | 0.47 | | | | | | 0.50 | | | |
| | Marginal $R^2$ / Conditional $R^2$ | | | 0.587 / 0.781 | | | | | | 0.615 / 0.809 | | | |

*Note.* All continuous predictors were centred. Degrees of freedom for *p*-values, standard errors and confidence intervals (CI) were computed using Satterthwaite's approximation. All *p*-values reported here are FDR-corrected and were computed using ANOVAs with type III sum of squares. Results that are significant on an alpha-level of 0.05 are marked with a star.

**Table S4. Results from models for control analysis (1-back vs. 2-back) of reading times for full original sample (N = 175).**

| | Control Analysis 2-back vs. 1-back: LMM for Full Original Sample (N = 175) | | | | | | |
|---|---|---|---|---|---|---|---|
| | ***Predictors*** | ***Estimate*** | ***Std. Error*** | ***CI*** | ***t*** | ***df*** | ***p*** |
| **Main Effects** | reading time of previous trial (log-transformed) | 0.062181 | 0.001755 | 0.058740 – 0.065622 | 35.420630 | 188475.92 | $3.296 \times 10^{-273}$ * |
| | d-prime | -0.006667 | 0.001898 | -0.010388 – -0.002947 | -3.512595 | 168020.99 | $7.398 \times 10^{-4}$ * |
| | mean d-prime singletasks | 0.094749 | 0.020772 | 0.053746 – 0.135752 | 4.561318 | 171.05 | $1.756 \times 10^{-5}$ * |
| | mean comprehension question performance | 0.003146 | 0.001352 | 0.000478 – 0.005815 | 2.327238 | 170.38 | $3.018 \times 10^{-2}$ * |
| | de-meaned comprehension question performance | 0.000038 | 0.000047 | -0.000053 – 0.00013 | 0.821932 | 150546.36 | $4.837 \times 10^{-1}$ |
| | word frequency | -0.128276 | 0.319365 | -0.756126 – 0.499574 | -0.401659 | 398.70 | $6.882 \times 10^{-1}$ |
| | word length | 0.005220 | 0.000414 | 0.004408 – 0.006031 | 12.616091 | 1351.55 | 4.020 × 10-34 * |
| | word entropy | 0.001711 | 0.000914 | -0.000081 – 0.003502 | 1.871897 | 4521.92 | $8.171 \times 10^{-2}$ |
| | n-back reaction [reaction vs. no reaction] | 0.316007 | 0.001917 | 0.312250 – 0.319764 | 164.836212 | 188099.24 | $< 1.33 \times 10^{-322}$ * |
| | block number | -0.011363 | 0.000295 | -0.011941 – -0.01079 | -38.535522 | 185049.49 | $1.33 \times 10^{-322}$ * |
| | trial number | -0.000450 | 0.000009 | -0.000467 – -0.000433 | -52.053811 | 10059.10 | $< 1.33 \times 10^{-322}$ * |
| | recording location [online vs. lab] | -0.226062 | 0.034131 | -0.293438 – -0.158686 | -6.623323 | 169.71 | $8.894 \times 10^{-10}$ * |
| | surprisal | 0.001848 | 0.000161 | 0.001532 – 0.002164 | 11.467327 | 2010.37 | $3.901 \times 10^{-29}$ * |
| | age | 0.008762 | 0.001101 | 0.006591 – 0.010934 | 7.961804 | 184.31 | $3.751 \times 10^{-13}$ * |
| | cognitive load [2-back vs. 1-back] | 0.338896 | 0.020892 | 0.297669 – 0.380123 | 16.221045 | 178.98 | $1.838 \times 10^{-36}$ * |
| **2-way Interactions** | surprisal x age | 0.000003 | 0.000005 | -0.000007 – 0.000012 | 0.545912 | 187940.82 | $6.159 \times 10^{-1}$ |
| | surprisal x cognitive load [2-back vs. 1-back] | -0.000148 | 0.000173 | -0.000486 – 0.000191 | -0.855146 | 187910.28 | $4.837 \times 10^{-1}$ |
| | age x cognitive load [2-back vs. 1-back] | 0.000689 | 0.001156 | -0.001593 – 0.00297 | 0.595977 | 172.03 | $6.133 \times 10^{-1}$ |
| **3-way Interaction** | surprisal x age x cognitive load [2-back vs. 1-back] | 0.000033 | 0.000010 | 0.000014 – 0.000052 | 3.372931 | 188203.53 | $1.144 \times 10^{-3}$ * |
| **Model Fit** | Intra-class Correlation (ICC) | | | 0.44 | | | |
| | Marginal $R^2$ / Conditional $R^2$ | | | 0.442 / 0.690 | | | |

*Note.* All continuous predictors were centred. Degrees of freedom for *p*-values, standard errors and confidence intervals (CI) were computed using Satterthwaite's approximation. All *p*-values reported here are FDR-corrected and were computed using ANOVAs with type III sum of squares. Results that are significant on an alpha-level of 0.05 are marked with a star.

**Table S5.** ***Results from GAM for control analysis of reading times for full original sample (N = 175).***

| Control Analysis: GAM for Full Original Sample (N = 175) | | | | | | | | |
|---|---|---|---|---|---|---|---|---|
| | ***Predictors*** | ***Estimate*** | ***Std. Error*** | ***t*** | ***F*** | ***EDF*** | ***p*** | |
| **Main Effects** | reading time of previous trial (log-transformed) | | | | 204.591 | 33.664 | $< 2 \times 10^{-16}$ | * |
| | d-prime | | | | 38.774 | 28.042 | $< 2 \times 10^{-16}$ | * |
| | mean d-prime singletasks | | | | 24.344 | 1.892 | $< 2 \times 10^{-16}$ | * |
| | mean comprehension question performance | | | | 5.348 | 2.305 | $3.23 \times 10^{-3}$ | * |
| | de-meaned comprehension question performance | | | | 39.408 | 7.477 | $< 2 \times 10^{-16}$ | * |
| | word frequency | | | | 6.837 | 7.571 | $< 2 \times 10^{-16}$ | * |
| | word length | | | | 73.076 | 4.038 | $< 2 \times 10^{-16}$ | * |
| | word entropy | | | | 4.027 | 3.704 | $2.44 \times 10^{-3}$ | * |
| | surprisal | | | | 9.547 | 4.107 | $< 2 \times 10^{-16}$ | * |
| | age | | | | 51.783 | 3.028 | $< 2 \times 10^{-16}$ | * |
| | n-back reaction [reaction vs. no reaction] | 0.3167 | 0.00179 | 177.06 | | | $< 2 \times 10^{-16}$ | * |
| | block number | -0.0061 | 0.00017 | -36.89 | | | $< 2 \times 10^{-16}$ | * |
| | trial number | -0.0004 | 0.00001 | -64.47 | | | $< 2 \times 10^{-16}$ | * |
| | recording location (online vs. lab) | -0.2514 | 0.02602 | -9.66 | | | $< 2 \times 10^{-16}$ | * |
| | cognitive load [1-back vs. Reading Only] | 0.4318 | 0.02514 | 17.17 | | | $< 2 \times 10^{-16}$ | * |
| | cognitive load [2-back vs. Reading Only] | 0.7819 | 0.02526 | 30.95 | | | $< 2 \times 10^{-16}$ | * |
| **2-way Interactions** | surprisal x cognitive load | | | | 13.962 | 13.849 | $< 2 \times 10^{-16}$ | * |
| **3-way Interactions** | surprisal x age x cognitive load [Reading Only] | | | | 23.946 | 10.248 | $< 2 \times 10^{-16}$ | * |
| | surprisal x age x cognitive load [1-back] | | | | 2.874 | 2.017 | $3.616 \times 10^{-2}$ | * |
| | surprisal x age x cognitive load [2-back] | | | | 2.392 | 4.877 | $2.375 \times 10^{-2}$ | * |
| **Random Effects** | Cognitive load \| ID | | | | 255.250 | 508.610 | $< 2 \times 10^{-16}$ | * |
| | Text Nr. | | | | 14053.220 | 7.840 | $< 2 \times 10^{-16}$ | * |
| | Word | | | | 1.870 | 804.600 | $< 2 \times 10^{-16}$ | * |
| | Colour | | | | 12.530 | 2.770 | $< 2 \times 10^{-16}$ | * |
| **Model Fit** | $R^2$ | | | | 815 | | | |

*Note. p*-values were computed using Wald's approximation as implemented in the package *mgcv*. Results that are significant on an alpha-level of 0.05 are marked with a star. EDF: Effective degrees of freedom.

**Table S6. Results from the model for reading times for full original sample (N = 175) for the effects of entropy, cognitive load and age on reading time.**

| | | LMM for Full Original Sample (N = 175) | | | | | |
|---|---|---|---|---|---|---|---|
| | ***Predictors*** | ***Estimate*** | ***Std. Error*** | ***CI*** | ***t*** | ***df*** | ***p*** |
| **Main Effects** | reading time of previous trial (log-transformed) | 0.110066 | 0.001410 | 0.107302 – 0.112830 | 78.048894 | 287686.846 | **< $8.542 \times 10^{-79}$** * |
| | d-prime | -0.006248 | 0.001767 | -0.009712 – -0.002784 | -3.535537 | 224708.066 | **$6.105 \times 10^{-4}$** * |
| | mean d-prime singletasks | 0.084628 | 0.019587 | 0.045963 – 0.123292 | 4.320683 | 169.899 | **$4.225 \times 10^{-5}$** * |
| | mean comprehension question performance | 0.002842 | 0.001276 | 0.000322 – 0.005362 | 2.226655 | 169.295 | **$3.275 \times 10^{-2}$** * |
| | de-meaned comprehension question performance | 0.000052 | 0.000039 | -0.000023 – 0.000127 | 1.350491 | 257990.129 | $1.769 \times 10^{-1}$ |
| | word frequency | 0.608299 | 0.360822 | -0.100080 – 1.316678 | 1.685873 | 725.615 | $9.929 \times 10^{-2}$ |
| | word length | 0.007812 | 0.000406 | 0.007015 – 0.008609 | 19.220944 | 1402.270 | **$9.864 \times 10^{-73}$** * |
| | surprisal | 0.001682 | 0.000150 | 0.001387 – 0.001977 | 11.177024 | 2359.191 | **$7.143 \times 10^{-28}$** * |
| | n-back reaction [reaction vs. no reaction] | 0.317360 | 0.001800 | 0.313832 – 0.320888 | 176.311387 | 287866.290 | **< $8.542 \times 10^{-79}$** * |
| | block number | -0.006481 | 0.000164 | -0.006803 – -0.006160 | -39.539875 | 287255.082 | **< $8.542 \times 10^{-79}$** * |
| | trial number | -0.000456 | 0.000007 | -0.000470 – -0.000442 | -64.065173 | 18557.632 | **< $8.542 \times 10^{-79}$** * |
| | recording location [online vs. lab] | -0.219370 | 0.032287 | -0.283107 – -0.155632 | -6.794462 | 168.720 | **$3.895 \times 10^{-10}$** * |
| | entropy | 0.001412 | 0.000785 | -0.000126 – 0.002950 | 1.800222 | 7570.659 | $8.213 \times 10^{-2}$ |
| | age | 0.009110 | 0.000991 | 0.007155 – 0.011065 | 9.195551 | 178.495 | **$2.325 \times 10^{-16}$** * |
| | cognitive load [1-back vs. Reading Only] | 0.473980 | 0.013924 | 0.446501 – 0.501458 | 34.041427 | 176.188 | **$8.542 \times 10^{-79}$** * |
| | cognitive load [2-back vs. Reading Only] | 0.791850 | 0.026098 | 0.740340 – 0.843361 | 30.341059 | 173.750 | **$7.294 \times 10^{-71}$** * |
| **2-way Interactions** | entropy x age | 0.000090 | 0.000030 | 0.000032 – 0.000148 | 3.030333 | 287391.572 | **$3.257 \times 10^{-3}$** * |
| | entropy x cognitive load [1-back vs. Reading Only] | 0.006638 | 0.001273 | 0.004142 – 0.009133 | 5.213908 | 287500.035 | **$3.416 \times 10^{-7}$** * |
| | entropy x cognitive load [2-back vs. Reading Only] | 0.006490 | 0.001291 | 0.003959 – 0.009021 | 5.025891 | 287757.942 | **$8.595 \times 10^{-7}$** * |
| | age x cognitive load [1-back vs. Reading Only] | -0.002785 | 0.000776 | -0.004317 – -0.001253 | -3.587447 | 171.994 | **$6.142 \times 10^{-4}$** * |
| | age x cognitive load [2-back vs. Reading Only] | -0.002441 | 0.001455 | -0.005313 – 0.000430 | -1.678106 | 170.772 | $9.929 \times 10^{-2}$ |
| **3-way Interactions** | entropy x age x cognitive load [1-back vs. Reading Only] | -0.000399 | 0.000072 | -0.000540 – -0.000258 | -5.546582 | 287440.357 | **$5.831 \times 10^{-8}$** * |
| | entropy x age x cognitive load [2-back vs. Reading Only] | -0.000188 | 0.000073 | -0.000331 – -0.000045 | -2.577310 | 287488.317 | **$1.258 \times 10^{-2}$** * |
| **Model Fit** | Intra-class Correlation (ICC) | | | 0.46 | | | |
| | Marginal $R^2$ / Conditional $R^2$ | | | 0.643 / 0.807 | | | |

*Note.* All continuous predictors were centred. Degrees of freedom for *p*-values, standard errors and confidence intervals (CI) were computed using Satterthwaite's approximation. All *p*-values reported here are FDR-corrected and were computed using ANOVAs with type III sum of squares. Results that are significant on an alpha-level of 0.05 are marked with a star.
